# Conjugation of dn-OPDA with amino acids inhibits its hormonal bioactivity in *Marchantia polymorpha*

**DOI:** 10.1101/2024.08.27.609933

**Authors:** Wenting Liang, Ángel M. Zamarreño, Salvador Torres-Montilla, Antonio de la Torre, Jean Chrisologue Totozafy, Takuya Kaji, Minoru Ueda, Massimiliano Corso, José M. García-Mina, Roberto Solano, Andrea Chini

**Author notes:** corresponding author’s.

## Abstract

Jasmonates are important phytohormones activating plant tolerance to biotic and abiotic stress, as well as different development processes. A conserved signalling pathway activated by distinct hormones in different plant species mediates these responses: dinor-12-oxo-phytodienoic acid (dn-OPDA) isomers in bryophytes and lycophytes, and JA-Ile in most vascular plants. The final responses depend, in many cases, on the accumulation of specialized metabolites. To identify novel compounds regulated by the dn-OPDA pathway in Marchantia, untargeted metabolomic analyses were carried out in response to dn-OPDA-regulated stress. A novel group of molecules were identified as dn-OPDA-amino acid conjugates (dn-OPDA-aas), and their accumulation after wounding and herbivory confirmed by targeted metabolic profiling in Marchantia and all species in which we previously found dn-*iso*-OPDA. Mutants in *GRETCHEN-HAGEN 3A* (Mp*GH3A*) failed to accumulate dn-OPDA-aa conjugates, and showed a constitutive activation of the OPDA pathway and increased resistance to herbivory. Our results show that dn-*iso*-OPDA bioactivity is reduced by conjugation with amino acids. Therefore, a dichotomous role of jasmonate conjugation in land plants is highlighted: jasmonic acid (JA) conjugation with isoleucine (Ile) produce the bioactive JA-Ile in tracheophytes, whereas conjugation of dn-*iso*-OPDA with different amino acids disactivate the hormone in bryophytes and lycophytes.

## Introduction

Jasmonates (JAs) are key regulators of plant development and adaptation to environmental stress, including defence against herbivores and necrotrophic pathogens (Howe et al., 2018; Gasperini and Howe 2024). Studies on *Arabidopsis thaliana*, defined the (+)-7-*iso*-JA-Ile as the ligand of the jasmonate co-receptor complex formed by the F-box coronatine insensitive 1 (COI1) and the jasmonate ZIM domain (JAZ) repressors (Xie et al., 1998; Chini et al., 2007; Thines et al., 2007; Katsir et al., 2008; Fonseca et al., 2009; Sheard et al., 2010). In contrast, studies in *Marchantia polymorpha* identified dn-*iso*-OPDA and Δ^4^-dn-*iso*-OPDA, collectively named here as dn-OPDAs, as the ligands of a conserved COI1/JAZ co-receptor (Monte et al., 2018; Kneeshaw et al., 2022). Recent studies showed that dn-OPDAs are the bioactive hormones in bryophytes and lycophytes, whereas JA-Ile is the hormone in most vascular plants, except some lycophytes (i.e., Lycopodiales) (Monte et al., 2022; Chini et al., 2023). Ligand binding specificity depends on a single amino acid in COI1 that alters the binding pocket size between vascular plants and bryophytes, and on specific residues within the JAZ degron of the JAZ co-receptors (Bowman et al., 2017; Monte et al., 2018; Monte et al., 2022).

The perception of the bioactive jasmonate induces the degradation of the JAZ repressors, which allows the activation of several transcription factors (TFs) regulating downstream jasmonate-dependent responses (Chini et al., 2007, 2009, 2016; Thines et al., 2007; Sheard et al., 2010; Fernández-Calvo et al., 2011; Monte et al., 2018, 2019; Peñuelas et al., 2019). Recently, an additional signalling role independent on COI1 has been described for the *cis* isomers of dn-OPDA and OPDA. These compounds are reactive electrophilic species (RES) that regulate thermotolerance, among other abiotic stresses, in different streptophyte plants, including the charophyte *Klebsormidium nitens*, the liverwort *M. polymorpha* or the angiosperm *Arabidopsis thaliana* (Farmer & Mueller, 2013; Monte et al., 2020).

The JAs biosynthetic pathway is fairly well understood. The octadecanoid and hexadecanoid pathways lead to the formation of OPDA and dn-OPDA, respectively, that are precursors of the bioactive JA-Ile in angiosperms (Wasternack, 2017; Chini et al., 2018; Yi et al., 2024). In contrast, in Marchantia MpFAD5 and MpDES6 are respectively required for the biosynthesis of the hexadecatrienoic acid (16:3) and eicosapentanoic acid (20:5), major precursors of the bioactive dn-OPDA and Δ^4^-dn-OPDA (Keenshaw et al., 2022; Soriano et al., 2022). In Arabidopsis, the final step in the biosynthesis of the bioactive JA-Ile is mediated by JAR1/GH3.11 and, to a lesser extent, by GH3.10 that conjugate JA to Ile (Staswick & Tiryaki, 2004; Delfin et al., 2022). The JAR1 enzyme and function is well conserved in several vascular plants (Jez, 2022; Chini et al., 2023). In addition, the conjugation of JA with amino acids other than Ile has been reported in several plants (Gutierrez et al., 2012). For example, JA-Ala, JA-Val, JA-Leu and JA-Met are natural molecules whose biosynthesis is induced by wounding in Arabidopsis, tomato and rice (Suza et al., 2010; Yan et al., 2016). These JA-aa molecules have been described as COI1-JAZ ligands with different affinity to the receptor compared to JA-Ile in angiosperm plants, such as Arabidopsis and tomato (Katsir et al., 2008; Fu et al., 2021).

GH3 enzymes can also catalyse the conjugation of amino acids with other phytohormones such as auxins and salicylic acid (SA) (Staswick et al., 2005; Westfall et al., 2012; Jez, 2022; Brunoni et al., 2020, 2023). IAA-aa conjugates were described as inactive and they were proposed as IAA storage forms (LeClere et al. 2002; Rampey et al. 2004). Plant *GH3* genes usually belong to a large gene family and their extensive functional redundancy hampers to define the precise role of individual enzymes (Gutierrez et al., 2012; Casanova-Sáez et al., 2022).

To identify novel compounds regulated by dn-OPDA in Marchantia, we carried out untargeted metabolomic analyses. dn-OPDA-aa conjugates were identified as an uncharacterised set of evolutionarily conserved dn-OPDA-regulated compounds. To study the biological role of these molecules, we studied the loss-of-function mutants of the Mp*GH3A* gene and found to be required for dn-OPDA conjugation with amino acids. Physiological, transcriptional and metabolic analyses revealed that Mp*gh3a^ge^* mutants showed a constitutively activated dn-OPDA pathway, suggesting that the activity of dn-*iso*-OPDA is diminished by its conjugation with amino acids.

## Results

### Untargeted metabolomics identify dn-OPDA-amino-acid conjugates in Marchantia polymorpha

To identify novel metabolites regulated by dn-OPDA in *M. polymorpha* we carried out untargeted metabolomics analyses of mock and wounded plants using liquid chromatography-mass spectrometry (LC-MS/MS) of crude extracts of WT (Tak-1) and Mp*coi1-2* mutant plants. In addition to the expected classes of jasmonate-regulated molecules previously identified in response to wounding in several plants, such as oxylipins, terpenoids, phenylpropanoids and flavonoids, we also identified a significant accumulation of a strikingly abundant class of unknown metabolites, putatively annotated as N-acyl amino acids derivates (Figure 1A). Further analysis of their putative structure based on MS/MS spectra revealed that some of these compounds could be isomeric forms of dn-OPDA conjugated to glutamic acid, histidine, glutamine and different methyl- or hydroxyl-derivates (Figure S1). These naturally occurring molecules had not been described to date. The wound-induced accumulation of putative dn-OPDA amino acid (dn-OPDA-aa) conjugates was slightly diminished in Mp*coi1-2* mutant, suggesting only a minor requirement of the COI1 pathway for their accumulation (Figure 1B).

**Figure 1.**
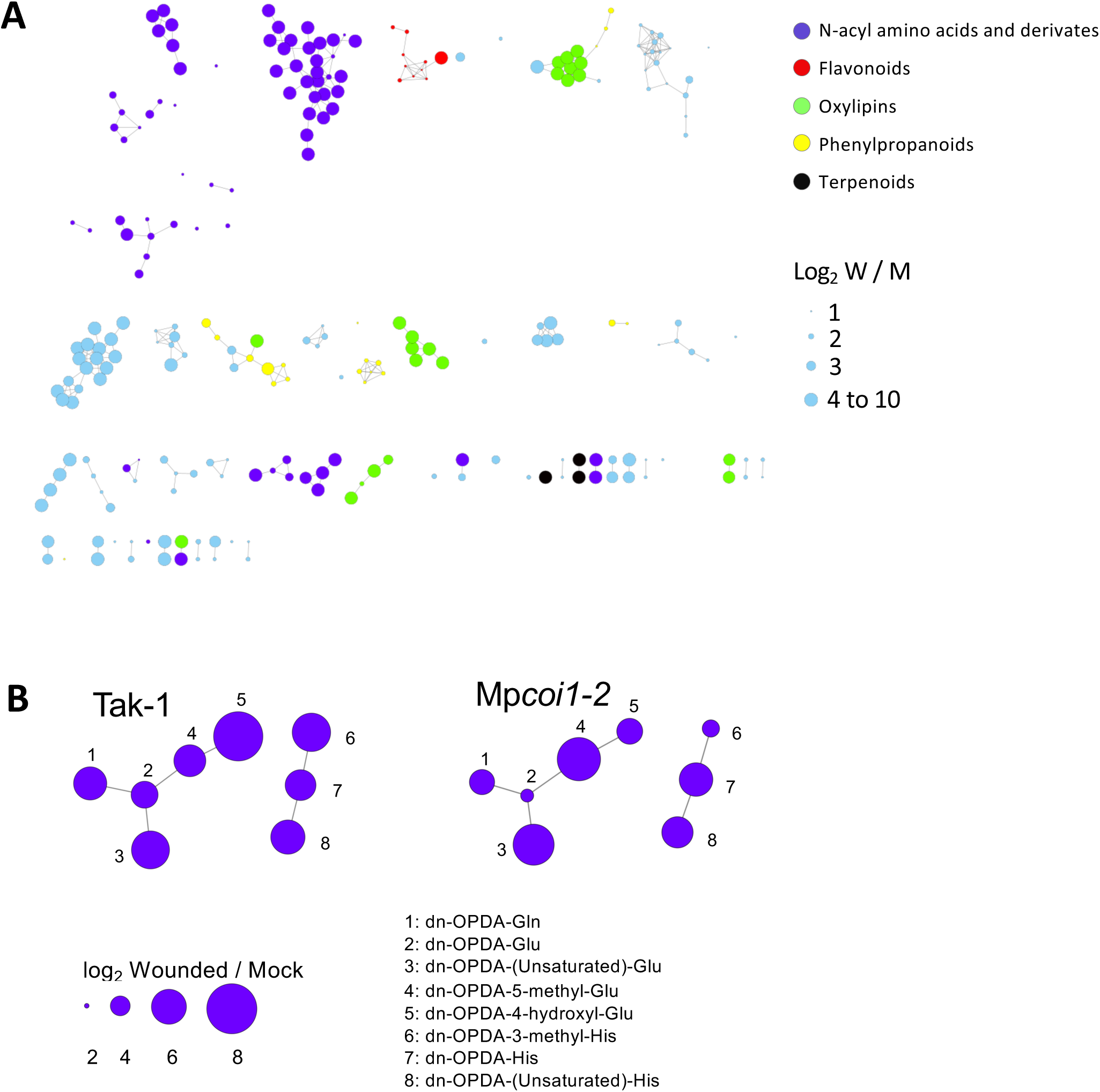
Molecular networks and annotation of Marchantia untargeted metabolomic profiles after wounding. (A) Molecular networks for metabolites differentially accumulated in wild-type (Tak-1) Marchantia after wounding. Analyses were carried out in positive ionization mode (ESI+) ionization modes. Metabolites are grouped based on their chemical structures and different colors correspond to different metabolic classes. The size of the nodes correlates with the accumulation promoted by wounding (log_2_ wound/mock). Cosine similarity scores of 0.7 were used for ESI+ analysis. (B) Comparison of molecular network for wound-induced putative dn-OPDA-aa conjugates class in wild-type (Tak-1) and Mp*coi1-2* mutant plants. The putative identity for each node is shown. The size of the nodes correlates with the induced accumulation upon wounding (log_2_ wound/mock).

To validate the ability of Marchantia plants to synthetise dn-OPDA-aa compounds after wounding, we analysed the accumulation of dn-OPDA conjugates with all natural amino acids in targeted LC-MS analyses. Nine dn-OPDA-aa conjugates were detected and most of them significantly accumulated in response to wounding (Figure S1 and S2). To confirm the identity of these dn-OPDA-aa compounds, we focused on the most abundant candidates and chemically synthetized pure dn-*iso*-OPDA-Glu, dn-*iso*-OPDA-Gln and dn-*iso*-OPDA-His to use them as standard in targeted LC-MS analyses. The subsequent analyses corroborated that these dn-*iso*-OPDA-aa conjugates significantly accumulate in Tak-1 after wounding (Figure 2A). In contrast, Mp*fad5* mutant, impaired in dn-*iso*-OPDA biosynthesis (Soriano et al., 2022), accumulated substantially less conjugates, than WT pants indicating that these compounds are primirly synthesized from the canonical dn-OPDA biosynthetic pathway (Figure 2A). Next, we analysed if another stress inducing dn-OPDA accumulation, such as herbivory, could also promote the accumulation of dn-*iso*-OPDA-aa conjugates. Indeed, Tak-1 plants accumulated high levels of dn-*iso*-OPDA-aa conjugates after herbivory challenge (Figure 2B). Similar to Mp*coi1*, Mp*mycy* mutant showed only minor defects in the accumulation of these compounds, indicating that the COI1/MYC pathway is only marginally involved in their biosynthesis (Figure 2B).

**Figure 2.**
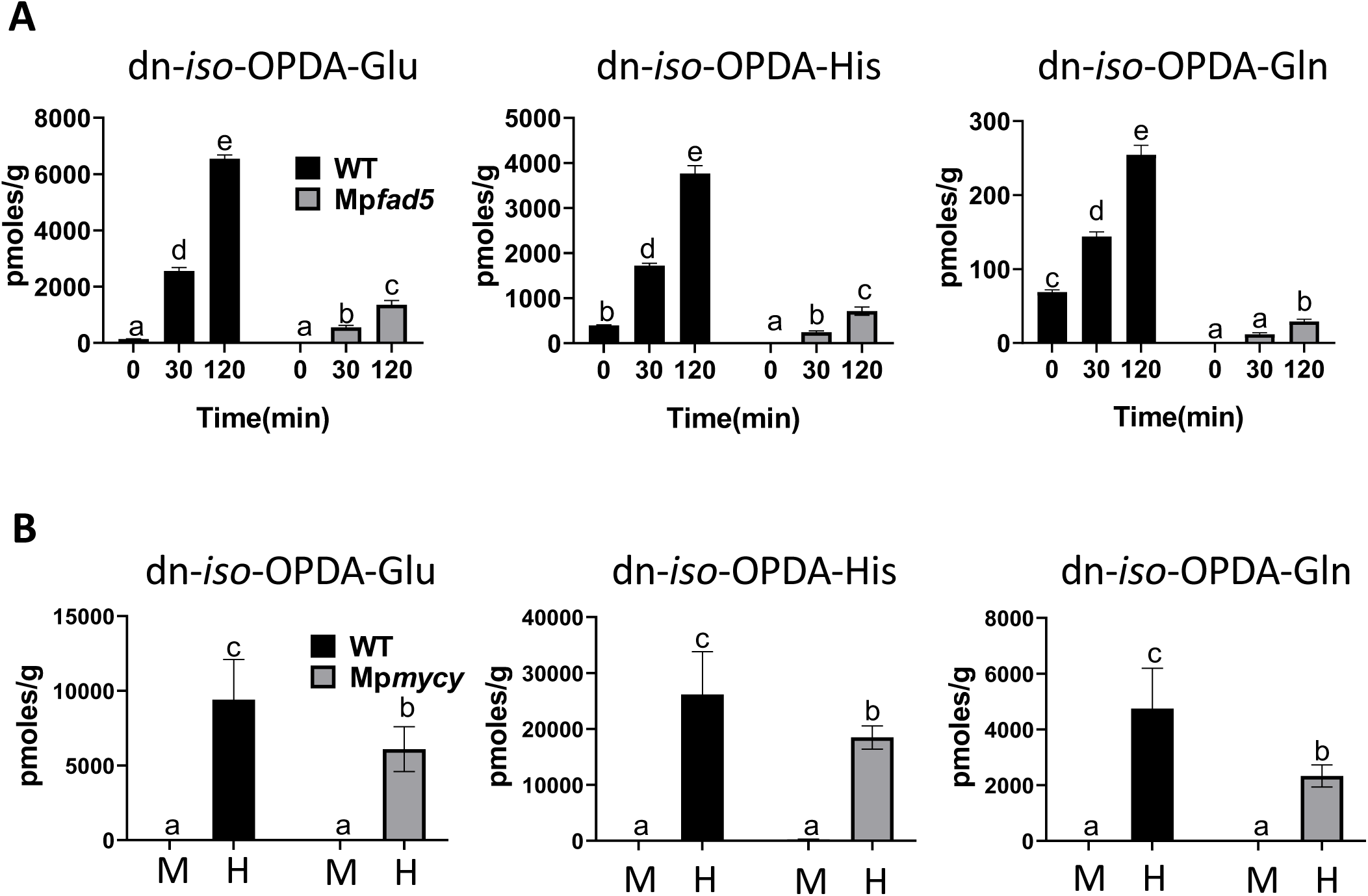
Accumulation of dn-*iso*-OPDA conjugates upon stress in *Marchantia polymorpha*. Time-course accumulation of dn-*iso*-OPDA conjugates [pmoles/fresh weight (g)] in wild-type (Tak-1) Marchantia plants, Mp*fad5* or Mp*mycy-1* mutants after wounding (A) or herbivory challenge (B). Plants were wounded and damaged tissues were collected after the indicated times (A). For herbivory assay, plants were challenged with *Spodoptera exigua* larvae and damaged tissues (H) were collected 8 days later (B). Unwounded (0) or not-challenged (mock, M) plants were included as control. Data shown as mean ± s.d. of three biological replicates. Experiments were repeated 3 times with similar results. Letters indicate significant different samples according to the one-way ANOVA/Tukey HSD post hoc test (P < 0.05).

In summary, metabolic profiling of Marchantia Tak-1 plants led to the identification of previously unknown derivates of the receptor-bioactive dn-*iso*-OPDA. The biosynthesis of dn-*iso*-OPDA-aa conjugates depended on the canonical dn-OPDA biosynthesis pathway (Soriano et al., 2020), but was mostly unaffected by the COI1 signalling pathway.

### The accumulation of dn-OPDA-amino acid conjugates is conserved in bryophytes and lycophytes

In addition to Marchantia, accumulation of dn-*iso*-OPDA has been recently reported in several bryophytes and lycophytes (Monte et al., 2022; Chini et al., 2023). To evaluate if dn-*iso*-OPDA-aa conjugates are synthetized in these plants, the accumulation of these compounds in response to wounding was studied in different lycophytes and bryophytes, as well as plants unable to accumulate dn-*iso*-OPDA, such as the angiosperm *Arabidopsis thaliana* and the charophyte *Klebsormidium nitens*.

As expected, in Arabidopsis and Klebsormidium, which lack dn-*iso*-OPDA, the corresponding dn-*iso*-OPDA-aa conjugates were not detected. In contrast, wounding induced the accumulation of dn-*iso*-OPDA-Glu and -Gln in the lycophyte *Huperzia Selago* and the bryophyte *Polytrichastrum formosum* (Figure 3), whereas the lycophyte *Selaginella lepidophylla* and the bryophytes *Physcomitrium patens* respectively accumulated dn-*iso*-OPDA-Glu and -Gln in response to wounding (Figure 3). dn-*iso*-OPDA-His was not detected in any of the analysed plants. Although the levels of and the ratio among dn-*iso*-OPDA-aa conjugates may vary in different plants, dn-*iso*-OPDA-Glu was the most accumulated among these compounds in all plants analysed with the only exception of *Selaginella lepidophylla*.

**Figure 3.**
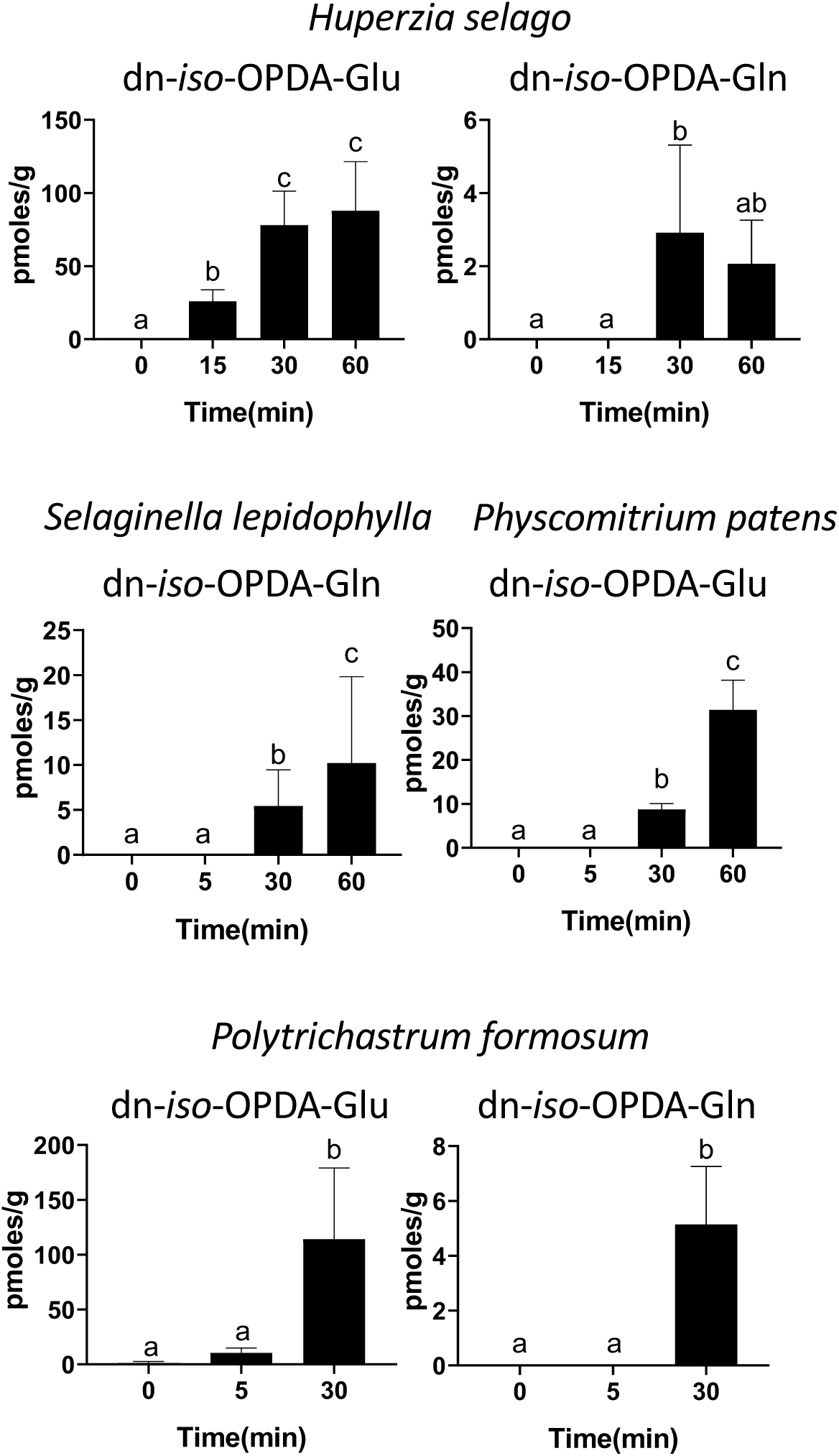
Accumulation of dn-*iso*-OPDA-aa conjugates in representative lycophytes and bryophytes. Time-course accumulation of dn-*iso*-OPDA conjugates [pmoles/fresh weight (g)] in lycophyte (*Selaginella lepidophylla* and *Huperzia selago*) and bryophyte (*Physcomitrum patens* and *Polytrichastrum formosum*) plants (N = 5-20). Plants were wounded and damaged tissues were collected after the indicated times. Unwounded plants (0) were included as control. Data shown as mean ± s.d. of three or four biological replicates. Experiments were repeated twice with similar results. Letters indicate significant different samples according to the one-way ANOVA/Tukey HSD post hoc test (P < 0.05).

Altogether, these data show that dn-OPDA-aa conjugates are not specifically occurring in Marchantia but they are evolutionarily conserved stress-inducible compounds in plants that synthesize dn-*iso*-OPDA as the bioactive hormone, including bryophytes and lycophytes.

### Role of MpGH3A in the biosynthesis of dn-OPDA-amino-acid conjugates

Conjugation of several phytohormones with amino acids have been reported (Jez, 2022). In the case of JAs, auxins and SA, the enzymes involved in amino-acid-conjugation belong to the GH3 family. Therefore, we reasoned that a GH3 enzyme may also conjugate dn-*iso*-OPDA with amino acids in Marchantia. The Marchantia genome contains two candidate *GH3* genes, Mp*GH3A* (Mp*6g07600*) and Mp*GH3B* (Mp*2g14010*). Mp*GH3A* was induced by OPDA and wounding in a MYC- and COI1-dependent manner, whereas Mp*GH3B* was not induced by OPDA (Figure S3). The fact that *JAR1*, encoding for the GH3 involved in jasmonate conjugation in Arabidopsis, is transcriptionally induced by jasmonate (Staswick & Tiryaki, 2004), pointed Mp*GH3A* as the most likely candidate to encode the enzyme mediating dn-OPDA conjugation with amino acids. To test this hypothesis, we generate loss-of-function mutants of Mp*GH3A* by CRISPR-Cas9.

First, two alleles were confirmed as genuine loss-of-function mutants. Mp*gh3a-1^ge^* carried a deletion of 569 nucleotide causing a substantial loss of the first exon and a premature stop codon (Figure S4A). The second allele Mp*gh3a-2^ge^* carried a deletion of 911 nucleotides, resulting in the loss of most of the third exon and a premature stop codon (Figure S4A); therefore, both alleles are predicted to be complete loss-of-function mutations. Next, the accumulation of dn-*iso*-OPDA-aa conjugates was analysed in Mp*gh3a-1^ge^* and Mp*gh3a-2^ge^*plants after wounding. Mp*gh3a^ge^* mutants were unable to accumulate dn-*iso*-OPDA-aa conjugates that were significantly induced by wounding in wild-type plants (Figure 4A). The only exception was the detection of traces of dn-*iso*-OPDA-Gln in Mp*gh3a^ge^*plants, although at significantly lower levels in wounded Mp*gh3a^ge^*plants compared to mock Tak-1 plants. In addition, dn-*cis*-OPDA and the bioactive dn-*iso*-OPDA and Δ^4^-dn-*iso*-OPDA accumulated more in Mp*gh3a ^ge^*mutants compared to Tak-1 plants after wounding (Figure 4B and S4B). In addition, Mp*gh3a^ge^* plants were significantly smaller than WT (Figure 4C-D). In summary, these results showed that MpGH3A is required *in planta* for the conjugation of dn-*iso*-OPDA with different amino acids.

**Figure 4.**
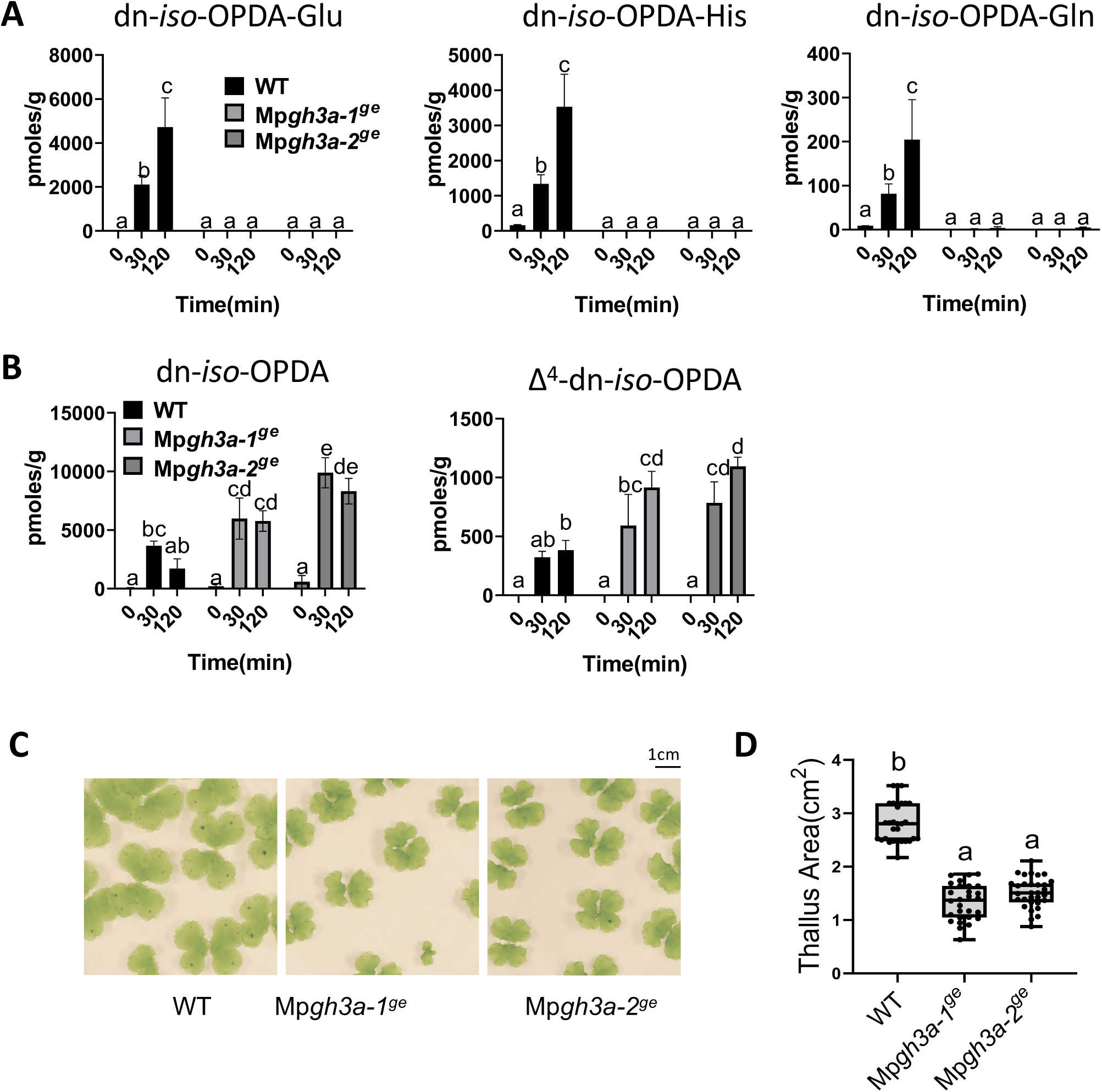
Accumulation of dn-*iso*-OPDAs in wild-type and Mp*gh3a^ge^* mutant plants after wounding. (A-B) Quantification of dn-*iso*-OPDAs and aa-conjugates levels [pmoles/fresh weight (g)] in Marchantia wild-type and Mp*gh3a^ge^* mutant plants after wounding. Plants were wounded, and damaged plants were collected after the indicated times. Data shown as mean ± s.d. of three biological replicates, 9 plants per each replicate. Experiments were repeated three times with similar results. Letters indicate significant different samples according to the one-way ANOVA/Tukey HSD post hoc test (P < 0.05). (C) Images of Marchantia wild-type and mutant plants grown for 21 days on GB5. Scale bars, 1 cm. (D) Quantification of thallus area (cm^2^) of plants (n = 30) shown in (C). The experiment was repeated three times with similar results. Box plots representation of thallus area; horizontal lines are medians, boxes show the upper and lower quartiles, and whiskers show the full data range.Letters indicate significant different samples according to the one-way ANOVA/Tukey HSD post hoc test (P < 0.05).

### Biological role of dn-OPDA-amino-acid conjugates in Marchantia polymorpha

In Arabidopsis, Ile conjugation with JA by AtGH3.11/JAR1 and AtGH3.10 produces the bioactive JA-Ile (Staswick & Tiryaki, 2004; Delfin et al., 2022). In contrast, conjugation of auxin and salicylic acid with different amino acids generate inactivate molecules. Therefore, amino acids conjugation can either activate or deactivate different signalling molecules.

To study the biological activity of the dn-*iso*-OPDA-aa conjugates in Marchantia, we analysed OPDA-regulated processes in Mp*gh3a ^ge^*plants. First, OPDA sensitivity of Mp*gh3a ^ge^* mutants was tested; OPDA induced significantly stronger growth inhibition in Mp*gh3a^ge^*mutants compared to Tak-1 plants (Figure 5), suggesting that Mp*gh3a ^ge^*plants response to OPDA is enhanced compared to WT. Next, analyses of dn-OPDA-specific marker genes showed that, the Mp*gh3a^ge^* plants showed significantly higher expression of Mp*DIR* and Mp*HBLH4* compared to the wild-type plants upon OPDA treatment (Figure S5). Finally, to evaluate the impact of the upregulation of dn-OPDA pathway on the transcriptome of Mp*gh3a-1^ge^*plants, RNA-Seq analyses were carried out on of mutants and WT plants were obtained. The Principal Component Analysis (PCA) plot showed that replicate samples clustered together, where treatment was the most severe factor affecting gene expression, with a lower impact of genotype (Supplemental Figure S6A). In basal conditions, 256 genes were differentially expressed (DEGs) in Mp*gh3a^ge^* compared to WT plants (148 up- and 108 down-regulated, Supplemental Table S1). Most of the up-regulated DEGs in Mp*gh3a-1^ge^* in basal conditions were induced by OPDA treatment in WT plants (Figure 6A), confirming a partial activation of the dn-OPDA pathway in the mutant. Gene ontology (GO) enrichment analysis showed that approximately half of the GO terms up-regulated in basal conditions in the Mp*gh3a-1^ge^* mutant are also up-regulated in response to OPDA treatment in WT plants (Supplemental Figure S6B and Supplemental Table S2). Furthermore, biological terms such as “response to JA”, “response to wounding” or response to additional stresses regulated by dn-OPDA were significantly up-regulated in Mp*gh3a-1^ge^* in basal conditions (Figure 6B and Supplemental Figure S6C). In response to OPDA, 142 genes showed a genotype:treatment interaction effect, responding to dn-OPDA in Mp*gh3a-1^ge^*differently to WT plants response (Supplemental Table S1). Most of these genes were up-regulated by OPDA in WT plants (Supplemental Figure S7A). In addition, weighted gene co-expression analysis (WGCNA) also defined different co-expressed modules; the vast majority of the genes (117) showed a steeper induction by OPDA in the mutant compared to WT (turquoise module; Figure 6C and Supplemental Figure 7B). In summary, the transcriptomic analyses showed that the absence of the *GH3A* gene enhances dn-OPDA-related gene expression both in basal conditions and after dn-OPDA treatment.

**Figure 5.**
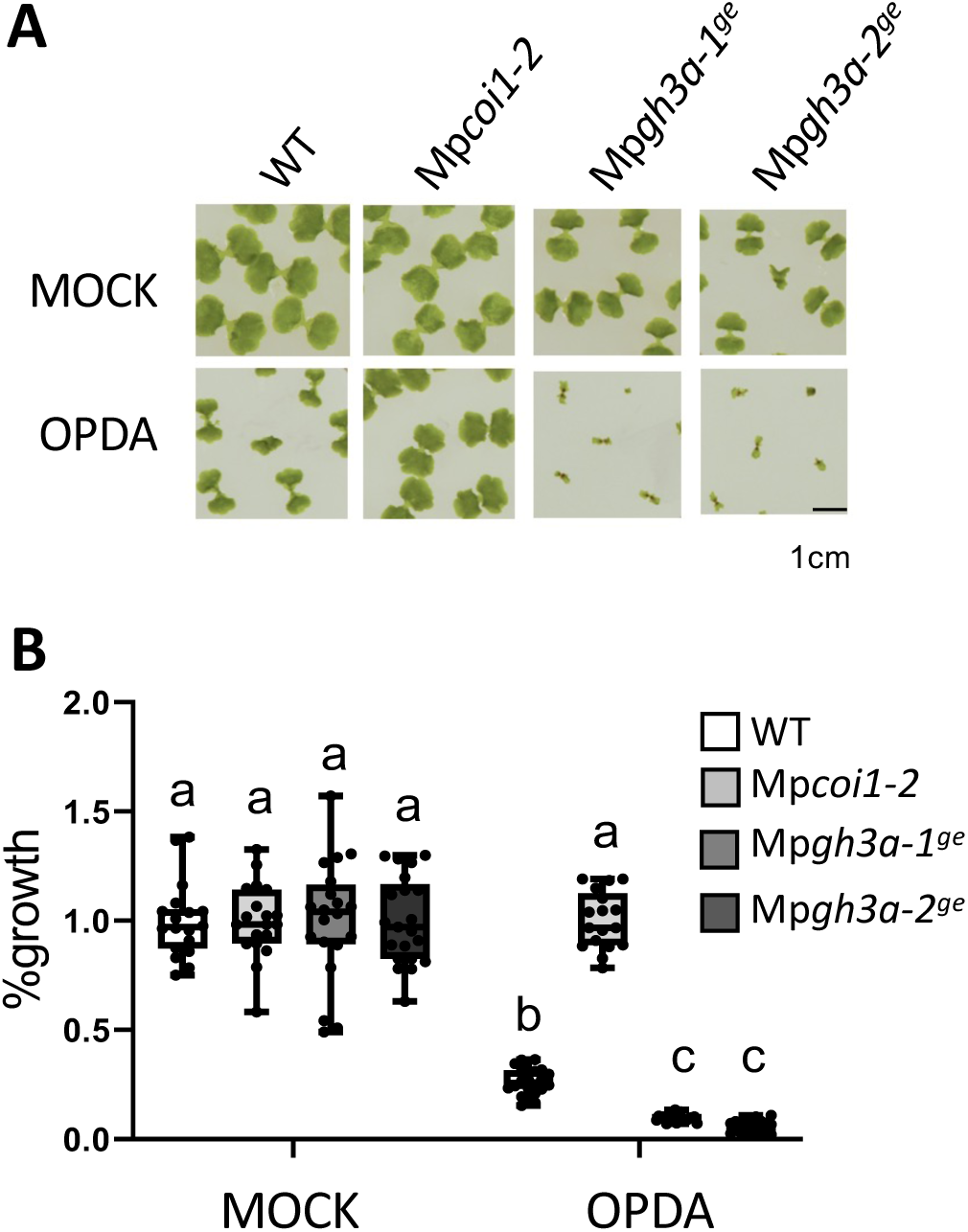
Effect of OPDA on the growth of wild-type and Mp*gh3a^ge^* mutant plants. (A) Images of the plants grown for 13 days in mock plates or in the presence of the dn-OPDA precursor OPDA. (B) Quantification of the growth inhibition induced by OPDA in wild-type, Mp*coi1-2* and Mp*gh3a^ge^* mutant plants grown for 13 days in absence (mock) and presence of 5 μM OPDA. Data shown as mean ± s.d. of three biological replicates, 20 plants per each replicate. Experiments were repeated 3 times with similar results. Box plots representation of growth inhibition. Letters indicate significant different samples according to the one-way ANOVA/Tukey HSD post hoc test (P < 0.05).

**Figure 6.**
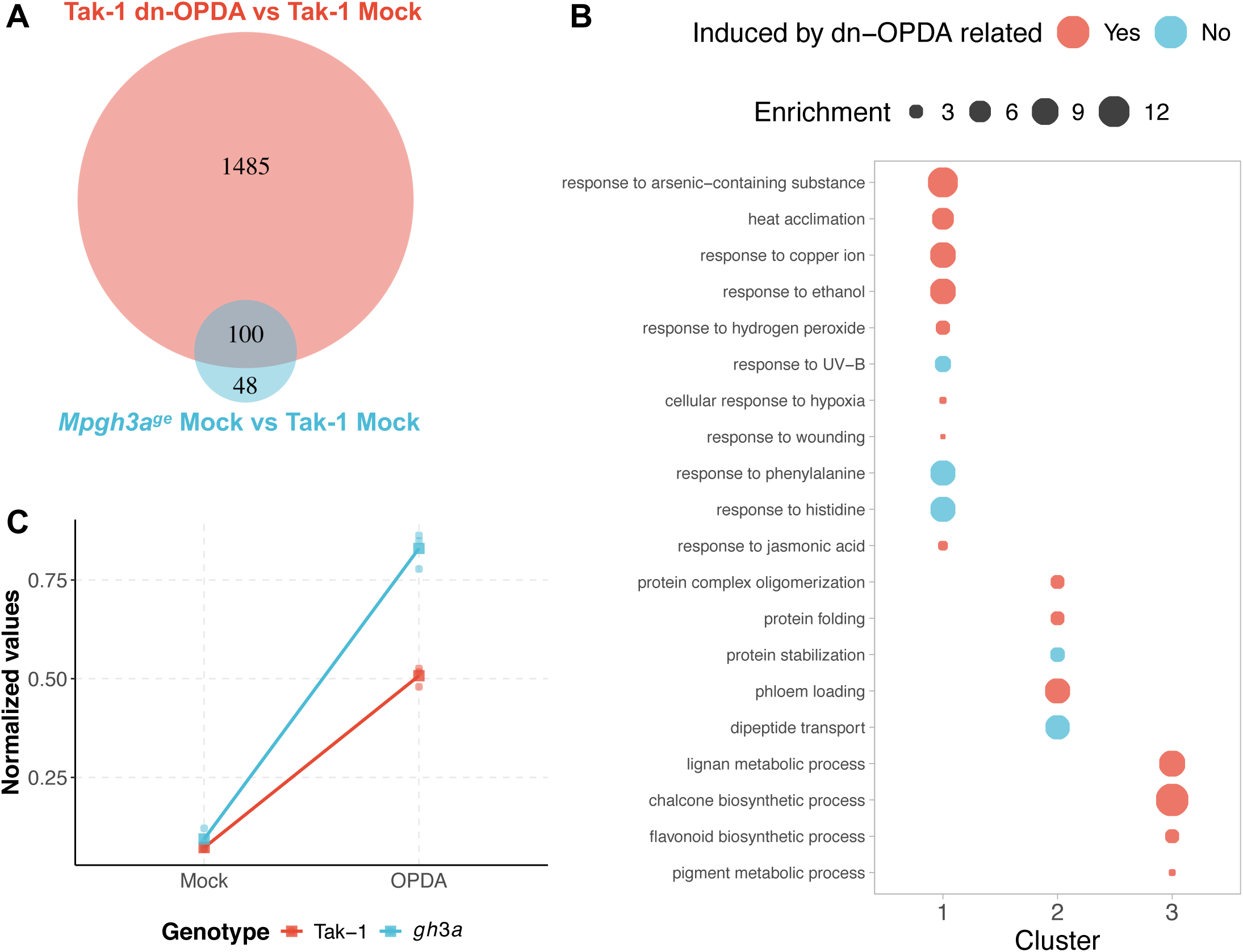
Transcriptomic analyses of *Mpgh3a-1^ge^*. (A) Venn diagram of differentially expressed genes (DEGs) up-regulated after dn-OPDA treatment in Tak1 (red) and DEGs up-regulated in *Mpgh3a-1^ge^*compared to Tak1 in Mock conditions (blue). (B) Top 20 Gene Ontology (GO) terms representing up-regulated DEGs in *Mpgh3a-1^ge^* compared to Tak1 in Mock conditions ordered according to the score integrating the p-value and the semantic algorithm. GO terms induced by OPDA treatment in WT pants are highlighted in red. The 3 GO clusters are shown in Figure S6. (C) Normalized expression of 117 genes (turquoise module) from the genotype:treatment interaction effect generated by Weighted Gene Co-expression Network Analysis (WGCNA). Green represents WT expression, while orange represents *Mpgh3a-1^ge^* expression.

Since the dn-OPDA pathway regulates biotic stress responses in Marchantia, including defences against the generalist herbivore Spodoptera (Monte et al., 2018; Peñuelas et al., 2019), we challenged Tak-1, Mp*gh3a^ge^*and Mp*coi1-2* plants with larvae of this insect. As previously reported, Mp*coi1-2* mutants were more susceptible than wild-type plants to herbivory. In contrast, Mp*gh3a ^ge^* mutants were significantly more resistant to *S. exigua* than WT plants (Figure 7).

**Figure 7.**
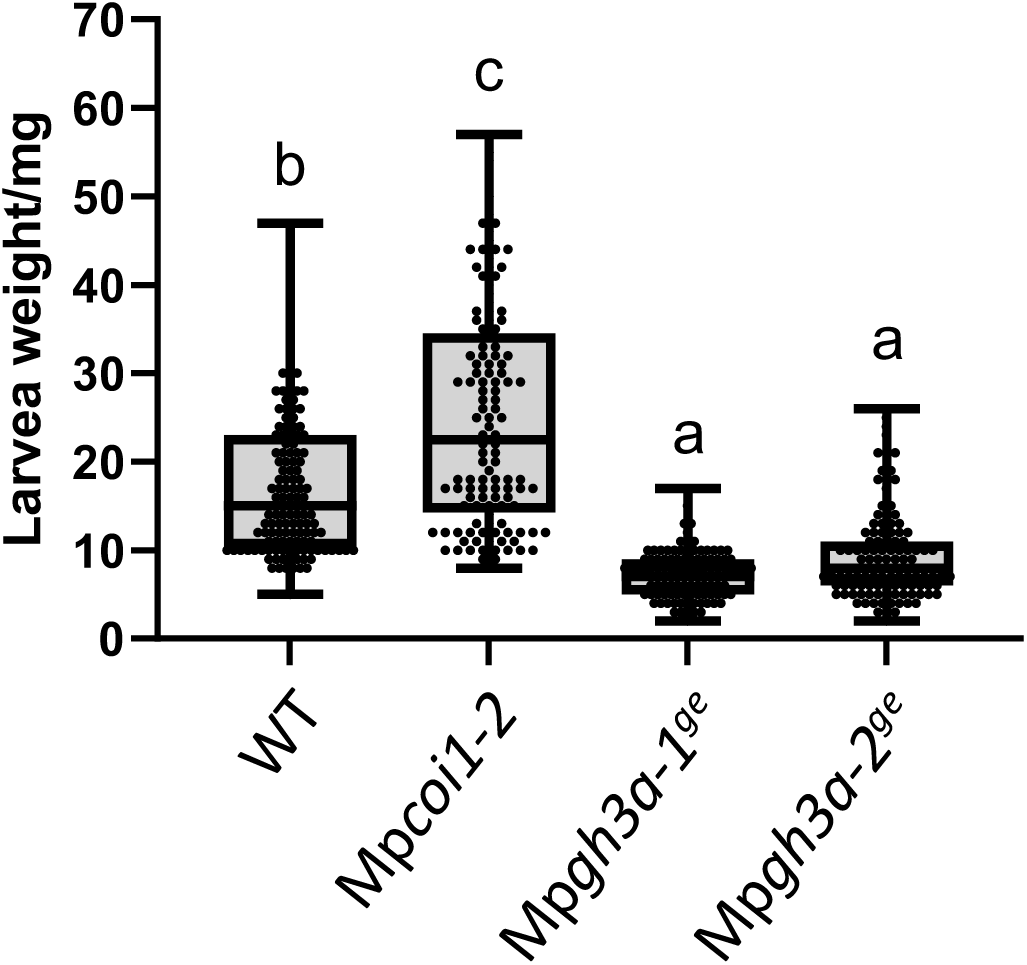
Mp*gh3a^ge^*mutants show enhanced resistance against *Spodoptera exigua*. *Spodoptera exigua* larval weight after 7 days feeding on Marchantia thalli of wild-type, Mp*coi1-2* and Mp*gh3a^ge^* mutants (n = 114-136). Box plots representation of larval weight. Different letters indicate significant differences evaluated by one-way ANOV A/Tukey HSD post hoc test (P < 0.05). Experiments were repeated four times with similar results.

Altogether, our results show that Mp*gh3a^ge^* mutants have an enhanced accumulation of dn-*iso*-OPDA, compared to wild type, due to the lack of amino acid conjugation. As a consequence, dn-*iso-*OPDA responses are significantly enhanced in the mutant than in WT plants. Therefore, these results show that conjugation with amino acids inactivates dn-*iso*-OPDA hormonal bioactivity in *M. polymorpha*.

## Discussion

Endogenous plant hormones, including jasmonates, are tightly regulated, since they modulate the vast majority of plant responses and developmental processes (Blázquez et al., 2020; Gasperini & Howe, 2024). Complementary to the biosynthetic pathway, hormone catabolism is a key process regulating the dynamic regulation of hormone homeostasis. For example, auxin homeostasis is regulated by several modifications of the bioactive indole-3-acetic acid (IAA), including oxidation, methylation and conjugation (Stepanova & Alonso, 2016; Abbas et al., 2018). It is well-described that GH3 enzymes catalyse the conjugation of amino acids with IAA, which in turn restrain its activity (LeClere et al. 2002; Rampey et al. 2004; Westfall et al., 2010; Jez, 2022). In contrast, Ile conjugation to JA produce the bioactive JA-Ile in tracheophyte plants, exemplifying that similar metabolic processes may have opposite biological effects (Staswick & Tiryaki, 2004; Chini et al., 2023).

Multiple processes have been reported to regulate jasmonates’ activity in angiosperms, including hydroxylation, carboxylation, decarboxylation, methylation, sulfation, and O-glycosylation (Wasternack & Strnad, 2018; Li et al., 2021; Yi et al., 2024). However, in Marchantia, and bryophytes in general, catabolic processes regulating the bioactive dn-OPDA homeostasis have not been reported to date.

Here, we describe an untargeted metabolomic analyses carried out in the bryophyte model *M. polymorpha* in response to wounding. In addition to compounds belonging to the canonical chemical families regulated by jasmonates in several plant species, such as oxylipins, terpenoids, phenylpropanoids and flavonoids, a novel family of compounds was identified (Figure 1). The synthesis of chemical standards and targeted LC-MS analyses confirmed these compounds as amino acid conjugates of dn-OPDA (Figure 2). To evaluated if conjugation of the bioactive dn-*iso*-OPDA regulates its activity, we first generated loss of function mutants of *MpGH3A* and showed that this enzyme is responsible for the synthesis of the conjugates (Figure 4). Besides the lack of dn-OPDA-aas, Mp*gh3a^ge^*mutants also showed a constitutive accumulation of the bioactive dn-OPDAs as well as activation of the dn-OPDA transcriptional cascade (Figure 4 and 6). Consistently, Mp*gh3a^ge^* plants showed OPDA hypersensitivity and enhanced dn-OPDA-regulated defences (Figure 5 and 7). Therefore, the analysis of Mp*gh3a^ge^*mutants suggested that dn-*iso*-OPDA-aa conjugates are inactive metabolites regulating the homeostasis of the bioactive dn-*iso*-OPDA.

In angiosperm plants, different set of amino acid conjugates to IAA have been reported as storage of the bioactive IAA, via regulated hydrolase, or irreversible inactive compounds (Rampey et al. 2004). The peptidases of the M20D family, including IAA-LEUCINE RESISTANT1 (ILR1) and its homologs, mediate the hydrolysis of auxin conjugates (Bartel and Fink 1995; LeClere et al. 2002). In contrast to auxin, salicylic acid conjugation to aspartic acid is irreversible since the SA-Asp conjugate cannot be converted into free SA (Chen et al., 2013). To date, the possibility that dn-*iso*-OPDA-aa conjugation is a reversable process is still unaddressed and future research will determine if amino acid conjugation is a regulated, reversible process. At this moment, we cannot rule out that, similar to IAA-aa conjugates (Kowalczyk and Sandberg, 2001), the most abundant dn-*iso*-OPDA-aa conjugates identified here may be irreversible inactive compounds, whereas less abundant dn-*iso*-OPDA-aa conjugates may be hydrolysed and their dynamic regulation may make it difficult to identify them *in planta*. Alternatively, similar to OPDA-aa conjugates, only a subgroup of dn-*iso*-OPDA-aa conjugates may undergo hydroxylation, independently on their accumulation levels (Brunoni et al., BioRxiv 2023; Siroka et al., accompanying manuscript). In this context, only one Marchantia homolog of angiosperm amidohydrolase was identified and *in vitro* analyses showed that MpILR can accept different auxin conjugates as substrates (Campanella et al., 2018; Bowman et al., 2021; Siroka et al., 2022). However, the naturally occurring IAA-Gly and IAA-Val were not among the MpILR substrates *in vitro* (Campanella et al., 2018). In addition, the amount of auxin conjugates in bryophytes is significantly lower than in angiosperms (Sztein et al. 1999; Záveská Drábková et al. 2015; Bowman et al., 2021). Indeed, under basal conditions, the levels of IAA conjugates were below our detection limits, similar to previous results (Kaneko et al., 2020), arguing against a role of MpILR in regulating auxin homeostasis. Therefore, to evaluate the hypothetical reversibility of inactive dn-*iso*-OPDA-aa conjugates to bioactive dn-*iso*-OPDA, it will be interesting to study the role of MpILR on dn-*iso*-OPDA-aa conjugates.

In contrast to Mp*GH3A*, Mp*GH3B* was not transcriptionally regulated by dn-OPDA (Monte et al., 2018; Peñuelas et al., 2019). However, the apparent lack of regulation of Mp*GH3B* by dn-OPDA does not rule out that MpGH3B may be involved in regulating the homeostasis of dn-*iso*-OPDA. For example, At*GH3.10* is weakly induced by JA, but AtGH3.10 enzyme could conjugate JA to several amino acids, showing a redundant role of AtGH3.10 and AtJAR1 in the biosynthesis of the bioactive JA-Ile in Arabidopsis (Delfin et al., 2022). Furthermore, although Mp*GH3A* expression is upregulated by dn-OPDA, the mutants in the perception and the signalling pathway of dn-OPDA, Mp*coi1* and Mp*mycy*, are only slightly affected in biosynthesis of dn-OPDA-aa conjugates (Figure 1 and 2), supporting a minor role of the transcriptional feedback loop in dn-OPDAs biosynthesis in Marchantia. These data also suggest that the basal level of MpGH3A enzyme is sufficient to promote the majority of the conjugation of dn-OPDA with amino acids. In this context, Mp*gh3a^ge^* mutant plants accumulated residual traces of dn-*iso*-OPDA-Gln after wounding (Figure 4), suggesting a minor redundancy in the dn-OPDA conjugation activity. Therefore, further research is required to define a putative role of MpGH3B in dn-OPDA homeostasis.

Furthermore, in the bryophyte and lycophyte plants analysed here, dn-*iso*-OPDA-Glu was the most abundant dn-*iso*-OPDA conjugate in most cases (Figure 3), whereas dn-*iso*-OPDA-His was only identified in *Marchantia polymorpha*. These results may suggest a species specificity in the biosynthesis of some of these compounds; however, due to the deactivation role of dn-*iso*-OPDA conjugates, it seems unlikely that different dn-*iso*-OPDA conjugates may hold specific function.

This work describes dn-*iso*-OPDA conjugation as the first process to inactivate the receptor-bioactive jasmonate molecule in Marchantia. However, additional processes regulating jasmonate homeostasis in angiosperms have been described, including hydroxylation, carboxylation, oxidation and sulfation among others (Wasternack & Strnad, 2018; Li et al., 2021). It will be interesting to evaluate if any additional jasmonate inactivation processes may be conserved in bryophytes. Noteworthy, in Arabidopsis different catabolic auxin processes, such as conjugation and oxidation, were described to be co-ordinately regulated to achive the most beneficial auxin homeostasis (Bowman et al., 2021). Therefore, it would be interesting to evaluate if additional dn-OPDA inactivation processes may interplay with dn-*iso*-OPDA-aa conjugation to obtain the best jasmonate homeostasis in bryophytes.

In summary, our results show that the signalling activity of dn-*iso*-OPDA is reduced by its conjugation with different amino acids. These results illustrate a dichotomous role of jasmonate conjugation in land plants: JA conjugation to Ile produces the bioactive JA-Ile in tracheophyte plants, whereas conjugation of dn-*iso*-OPDA to amino acids disactivate the receptor-bioactive dn-*iso*-OPDA in bryophyte and lycophyte plants.

## Material and Methods

### Plant material

*Marchantia polymorpha* accession Takaragaike-1 (Tak-1) was used as the wild-type (WT). In this genetic background, we used CRISPR-Cas9 nickase-mediated mutagenesis with Mp*GH3A* as the target. Four different pairs of gRNAs were designed in the first exon of the gene (Table S1). The eight gRNAs were then first cloned into pBC-GE12, pBC-GE23, pBC-GE34 and pMPGE_EN04 vectors. Next, they were transferred by LR reaction into the pMpGE017 binary vector carrying the CRISPRCas9 nickase. Wild-type plants were transformed and thalli selected by hygromycin resistance employing the regenerating thalli transformation (Kubota et al., 2013). To select mutant plants, genomic DNA of hygromycin-resistant explants were extracted and sequenced as previously described (Kneeshaw et al., 2022; Soriano et al., 2022). In addition, the Mp*coi1-2* and Mp*mycy* mutants were used as control in some experiments (Monte et al., 2018; Peñuelas et al., 2019).

### Chemicals

*cis*-12-oxo-phytodienoic acid (OPDA) and dinor-12-oxophytodienoic acid (dn-OPDA) were purchased from Cayman Chemical Co. (Ann Arbor, MI, USA), whereas dn-*cis*-OPDA and dn-*iso*-OPDA were previously synthesized (Chini et al., 2023).

### Culture conditions and wounding

Plants were routinely grown at 21°C, under continuous white light (50-60 mmol m^-2^ s^-1^) on half-strength Gamborg’s B5 medium Petri plates containing 1% agar. For jasmonate quantification, plants were grown for 2 weeks. For wounding assays, thalli of the WT and mutant plants were mechanically wounded with tweezers, pressing all over the thallus surface (Kneeshaw et al., 2022; Soriano et al., 2022). Wounded thalli were harvested and immediately frozen in liquid nitrogen after the described times.

### Growth inhibition assays

Growth inhibition assays were performed as previously described (Kneeshaw et al., 2022; Soriano et al., 2022). Briefly, OPDA was added to the GB5 media at a 5 μM concentration. After 2 weeks, pictures of the plants were acquired with a NIKON D1-x digital camera and the area of each plant was estimated with the ImageJ software. The growth percentage was calculated as the ratio of the area of hormone-treated versus untreated plants. The assays were repeated at least 3 times with similar results.

### Extraction of polar and semi-polar specialized metabolites

Polar and semi-polar metabolite fraction was extracted as previously described (Routaboul et al., 2006), with some modifications. Samples consisted of 5 mg of *M. polymorpha* grinded tissues. Briefly, 1 mL of Methanol/Water (80/20) + 0.05% of formic acid were added to each sample. The mixtures were sonicated using an ice-cooled ultrasonication (Fischer Scientific FB15050) for 1 min, then shaken in 2-ml tubes using a ThermoMixer™ C (Eppendorf) at 1400 rpm for 30 min and 4°C. The tubes were centrifuged at 10 000 rpm for 10 min at 4°C; the supernatants were collected in glass tubes. Next, 1 mL of extraction solution were added to the 2ml tube containing the pellet, and the extraction procedure was repeated. The two supernatants were pooled and dried using a vacuum concentrator (SpeedVac). The dried extracts were resuspended in 100 µL Acetonitrile:Water (1:9) of ULC/MS grade. The resuspended extracts were filtered with a glass microfiber filters (Cat. NO. 1820-037, Whatman International Ltd., UK) and distributed in HPLC vials. A Quality Control (QC) made up of 5 µL from each sample was prepared in a separate HPLC vials.

### UPLC-MS/MS analysis of polar and semi-polar specialized metabolites and UPLC-MS/MS data processing

Extracted samples were analysed as previously described (Boutet et al., 2022). The untargeted metabolomic data were acquired using a UHPLC system (Ultimate 3000 Thermo) coupled to quadrupole time of flight mass spectrometer (Q-TOF Impact II Bruker Daltonics, Bremen, Germany). A Nucleoshell RP 18 plus reversed-phase column (2x 100 mm, 2.7 µm; Macherey-Nagel) was used for chromatographic separation with the mobile phases consisting in: (A) 0.1% formic acid in water; and (B) 0.1% formic acid in acetonitrile. The flow rate was 400 µL min-1, and the following gradient was used: 95% (A) for 1 min, followed by a linear gradient from 95% (A) to 80% (A) from 1 to 3 min, then a linear gradient from 80% (A) to 75% (A) from 3 to 8 min, a linear gradient from 75% (A) to 40% (A) from 8 to 20 min; then, 0% of (A) was held until 24 min, followed by a linear gradient from 0% (A) to 95% (A) from 24 to 27 min. Finally, the column was washed by 30% (A) at for 3.5 min then re-equilibrated for 3.5 min (35 min total run time). Mass spectrometer data were obtained with data-dependent acquisition (DDA) method in both positive and negative electrospray ionisation (ESI) modes using the following parameters: capillary voltage, 4.5 kV; nebulizer gas flow, 2.1 bar; dry gas flow, 6.L.min-1; drying gas in the heated electrospray source temperature, 200°C. Samples were analysed at 8 Hz with a mass range of 100–1500 m/z. Stepping acquisition parameters were created to improve the fragmentation profile with a collision RF from 200 to 700 Vpp, a transfer time from 20 to 70 µsec, and collision energy from 20 to 40 eV. Each cycle included a MS fullscan and 5 MS/MS CID on the 5 main ions of the previous MS spectrum.

The UPLC-MS/MS data processing pipeline was performed as in (Boutet et al., 2022) with modifications. Briefly, .d data files (Bruker Daltonics, Bremen, Germany) were converted to .mzXML format using the MSConvert software (ProteoWizard package 3.0). mzXML data processing, mass detection, chromatogram building, deconvolution, samples alignment and data export were performed using MZmine 2 software (http://mzmine.github.io/) for both positive and negative data files. The ADAP chromatogram builder method was used with a minimum group size of scan 3, a group intensity threshold of 1000, a minimum highest intensity of 1000 and m/z tolerance of 10 ppm. Deconvolution was performed with the ADAP wavelets algorithm using the following setting: S/N threshold 8, peak duration range = 0.01–2 min RT wavelet range 0.01–0.2 min, MS2 scan were paired using a m/z tolerance range of 0.05 Da and RT tolerance of 0.2 min. Then, isotopic peak grouper algorithm was used with a m/z tolerance of 10 ppm and RT tolerance of 0.1. All the peaks were filtered using feature list row filter keeping only peaks with MS2 scan. The alignment of samples was performed using the join aligner with an m/z tolerance of 10 ppm, a weight for m/z and RT at 1, a retention time tolerance of 0.2 min. Metabolites accumulation was normalized according to the weight of seeds used for the extraction. Pooled QC sample injections across LC-MS/MS run were used to evaluate the quality of the run and untargeted metabolomic dataset. The relative standard deviation across the QC samples was calculated for each metabolic feature detected. The metabolic features were filtered according to their QC variation and only metabolic features with QC variation < 25% were kept.

### Molecular network and annotation of untargeted metabolomic data

As described previously, molecular networks were generated with MetGem software 20(https://metgem.github.io) using the .mgf and .csv files obtained with MZmine2 analysis. The molecular network was optimized for the ESI+ and ESI-datasets, and different cosine similarity score thresholds were tested. ESI - and ESI+ molecular networks were both generated using a cosine score threshold of 0.7 Molecular networks were exported to Cytoscape software (https://cytoscape.org/) to format the metabolic categories.

Metabolite annotation was performed in four consecutive steps. First, the ‘custom database search’ module from Mzmine was used to compare the obtained LC-MS/MS data with the IJPB chemistry and metabolomic platform homemade experimental (m/z absolute tolerance of 0.005 and RT tolerance of 0.2 min) and exact mass (m/z absolute tolerance of 0.005 Da or 10 ppm) libraries containing respectively 166 standards or experimental common features (RT, m/z) and 1112 ion known m/z. Second, the ESI – and ESI+ metabolomic data used for molecular network analyses were searched against the available MS2 spectral libraries (Massbank NA, GNPS Public Spectral Library, NIST14 Tandem, NIH Natural Product and MS-Dial), with absolute m/z tolerance of 0.02, 4 minimum matched peaks and minimal cosine score of 0.8 (Fig. 3a). Third, not-annotated metabolites that belong to molecular network clusters containing annotated metabolites from steps 1 and 2 were assigned to the same metabolic category. Following this annotation approach, 20 % of the metabolic features of the LC-MS/MS dataset were assigned to one of the metabolic categories identified. Fourth, the putative annotations of key metabolic features that were highlighted by statistical analyses was verified using Sirius, software or by checking the M2 spectra.

### Metabolites table and redundancy in mass spectrometry

Redundancy cleaning due to the mass spectrometry technique was performed using a local application (see Code availability) built using the freely available Shiny R package (https://cran.r-project.org/web/packages/shiny/index.html). It allows to identify the formation of adducts complementary to the protonated form in positive mode as: sodium potassium and ammonium and de-protonated in negative mode as: sodium, potassium, chlorine, formic acid, nitric acid, acetate, formic acid clusters associated with sodium by comparing the masses and retention time.

In the same way, the complementarity of positive and negative modes for certain metabolites was also eliminated by cross-referencing the modes of these complementary forms with those of their adducted forms. Both analyses were done using a mass tolerance parameter of 0.006 Da and a retention time of 0.2 minutes.

### Statistical analyses of metabolomic data

Statistical analyses were performed using Metaboanalyst 5.0 software (Pang et al., 2021). In particular, an ANOVA has been conducted to identify differentially accumulated metabolites among control and stress condition, and genotypes (adjusted p value < 0.05).

### Synthesis of dn-*iso*-OPDA amino acid conjugates

#### Synthesis of dinor-iso-OPDA-Glu

To a solution of dinor-*iso*-OPDA (Wang, Sakurai et al., 2021) (10.8 mg, 41 µmol, 1 eq.) and Et_3_N (12.0 µL, 82 µmol, 2 eq.) in THF (0.40 mL) was added ethyl chloroformate (4.0 µL, 41 µmol, 1 eq.) at 0 ℃. After being stirred at 0 ℃ for 2 h, sodium L-glutamate monohydrate (15.3 mg, 82 µmol, 2 eq.) dissolved in aq. diisopropylethylamine (45 µL, 246 µmol, 6 eq.) in H_2_O (400 µL) was added the above mixture and stirred for 2 h at room temperature. The reaction mixture was then acidified with 1M aq. HCl and extracted with CHCl_3_. The combined organic layer was washed with brine, dried over Na_2_SO_4_, and concentrated under reduced pressure. The residue was directly purified by RP-HPLC after membrane filtration (COSMOSIL^®^ Cholester column F20×250 mm, Flow rate 8.0 mL/min, detection 220 nm, eluent: MeCN+1%AcOH, B: H_2_O+1%AcOH, 0-5 min 30%A, 5-20 min 30-70%A, 20-25min 70%A, 25-26 min 70-100%A, 26-30 min 100%A; t_R_ 20.6 min) to give dinor-*iso*-OPDA-Glu (7.3 mg, 45%) as a pale yellow oil (Supplemental Figure S8). [a] ^19^ −7.33 (c 0.35, MeOH); ^1^H (400 MHz, CD_3_OD) d 5.26 (dt, *J* = 10.5, 7.3 Hz, 1H), 5.09 (dt, *J* = 10.5, 7.3 Hz, 1H), 4.33 (dd, *J* = 9.4, 5.0 Hz, 1H), 2.83 (d, *J* = 6.9 Hz, 2H), 2.50-2.46 (m, 2H), 2.41 (t, *J* = 7.3 Hz, 2H), 2.33-2.23 (m, 4H), 2.17 (t, *J* = 7.3 Hz, 2H), 2.13-2.02 (m, 1H), 2.07 (quin, *J* = 7.3 Hz, 2H), 1.84 (sext, *J* = 7.3 Hz, 1H), 1.57 (quin, *J* = 7.3 Hz, 2H), 1.50 (quin, *J* = 7.3 Hz, 2H), 1.30 (quin, *J* = 7.3 Hz, 2H), 0.86 (t, *J* = 7.3 Hz, 3H); ^13^C (100 MHz, CD_3_OD) d 212.44, 178.07, 176.27, 176.20, 174.95, 140.01, 133.19, 126.44, 52.94, 36.53, 35.15, 32.17, 31.26, 30.25, 30.19, 28.18, 27.86, 26.61, 21.93, 21.54, 14.54: HRMS (ESI neg.) [M-H]^-^ C_21_H_30_NO_6_^-^, calcd. 392.2079, found 392.2068

#### dinor-iso-OPDA-Gln

To a solution of dinor-*iso*-OPDA (10.8 mg, 41 µmol, 1 eq.) and Et_3_N (12.0 µL, 82 µmol, 2 eq.) in THF (0.40 mL) was added ethyl chloroformate (4.0 µL, 41 µmol, 1 eq.) at 0 ℃. After being stirred at 0 ℃ for 2 h, L-glutamine (12.1 mg, 82 µmol, 2 eq.) dissolved in aq. diisopropylethylamine (45 µL, 246 µmol, 6 eq.) in H_2_O (400 µL) was added the above mixture and stirred for 2 h at room temperature. The reaction mixture was then acidified with 1M aq. HCl and extracted with CHCl_3_. The combined organic layer was washed with brine, dried over Na_2_SO_4_, and concentrated under reduced pressure. The residue was directly purified by RP-HPLC after membrane filtration (COSMOSIL^®^ Cholester column F20×250 mm, Flow rate 8.0 mL/min, detection 220 nm, eluent: MeCN+1%AcOH, B: H_2_O+1%AcOH, 0-5 min 30%A, 5-20 min 30-70%A, 20-25min 70%A, 25-26 min 70-100%A, 26-30 min 100%A; t_R_ 18.6 min) to give dinor-iso-OPDA-Glu (5.3 mg, 33%) as a pale yellow oil (Supplemental Figure S8). [a] ^18^ −2.78 (c 0.27, MeOH); ^1^H (400 MHz, CD_3_OD) d 5.26 (dtt, *J* = 10.5, 7.3, 1.4 Hz, 1H), 5.09 (dtt, *J* = 10.5, 7.3, 1.4 Hz, 1H), 4.29 (dd, *J* = 9.2, 5.0 Hz, 1H), 2.83 (d, *J* = 7.3 Hz, 2H), 2.51-2.45 (m, 2H), 2.41 (t, *J* = 7.8 Hz, 2H), 2.29-2.13 (m, 7H), 2.06 (quin, *J* = 7.3 Hz, 2H), 1.84 (sext, *J* = 7.3 Hz, 1H), 1.57 (m, 2H), 1.50 (quin, *J* = 7.3 Hz, 2H), 1.30 (quin, *J* = 7.3 Hz, 2H), 0.89 (t, *J* = 7.3 Hz, 3H); ^13^C (100 MHz, CD_3_OD) d 212.43, 178.07, 177.67, 176.20, 174.87, 140.01, 133.20, 126.44, 53.21, 36.56, 35.15, 32.77, 32.17, 30.27, 30.19, 28.51, 28.18, 26.59, 21.93, 21.54, 14.54: HRMS (ESI neg.) [M-H]^-^C_21_H_31_N_2_O_5_^-^, calcd. 391.2238, found 391.2233

#### dinor-iso-OPDA-His

To a solution of dinor-*iso*-OPDA (10.8 mg, 41 µmol, 1 eq.) and Et_3_N (12.0 µL, 82 µmol, 2 eq.) in THF (0.40 mL) was added ethyl chloroformate (4.0 µL, 41 µmol, 1 eq.) at 0 ℃. After being stirred at 0 ℃ for 2 h, L-Histidine (12.8 mg, 82 µmol, 2 eq.) dissolved in aq. diisopropylethylamine (45 µL, 246 µmol, 6 eq.) in H_2_O (400 µL) was added the above mixture and stirred for 2 h at room temperature. The reaction mixture was then acidified with 1M aq. HCl and washed with CHCl_3_. The remaining aqueous layer was concentrated under reduced pressure. The residue was directly purified by RP-HPLC after membrane filtration (COSMOSIL^®^ Cholester column F20×250 mm, Flow rate 8.0 mL/min, detection 220 nm, eluent: MeCN+1%AcOH, B: H_2_O+1%AcOH, 0-5 min 30%A, 5-20 min 30-50%A, 20-21min 50-100%A, 21-25 min 100%A; t_R_ 14.4 min) to give dinor-iso-OPDA-His (4.8 mg, 29%) as a pale yellow oil (Supplemental Figure S8). [a] ^19^ −4.81 (c 0.24, MeOH); ^1^H (400 MHz, CD OD) d 8.83 (s, 1H), 7.33 (s, 1H), 5.34 (dt, *J* = 10.5, 7.3 Hz, 1H), 5.17 (dt, *J* = 10.5, 7.3 Hz, 1H), 4.77 (dd, *J* = 8.7, 5.0 Hz, 1H), 3.34-3.25 (m, 1H), 3.09 (dd, *J* = 15.6, 9.2 Hz, 1H), 2.91(d, *J* = 6.9 Hz, 2H), 2.56 (m, 2H), 2.48 (t, *J* = 7.3 Hz, 2H), 2.35 (m, 2H), 2.28-2.14 (m, 2H), 2.15 (quint, *J* = 7.3 Hz, 2H), 1.64-1.50 (m, 4H), 1.40-1.30 (m, 2H), 0.97 (t, *J* = 7.3 Hz, 3H); ^13^C (100 MHz, CD_3_OD) d 212.34, 177.88, 176.02, 173.12, 140.02, 135.09, 133.19, 131.43, 126.43, 118.38, 52.41, 36.50, 35.14, 32.11, 30.21, 30.19, 28.14 (2C), 26.53, 21.93, 21.53, 14.53: HRMS (ESI neg.) [M-H]^-^ C_22_H_30_N_3_O_4_^-^, calcd. 400.2242, found 400.2234.

#### Chemical compound measurements

Measurement of the described compounds was carried out as previously described (Chini et al., 2018; Chini et al., 2023). Tissue from 5 to 10 plants was pooled for each biological sample and at least three independent biological replicates were performed for each treatment. Briefly, endogenous dn-OPDAs and derivate compounds were analysed using high-performance liquid chromatography electrospray-high resolution accurate MS (HPLC-ESI-HRMS) using an Orbitrap Exploris 120 detector in Full Scan and Product Ion Scan modes. The experiment was repeated twice to four times with similar results. Data are shown as mean ± SD.

#### Transcriptional analyses

For gene expression analyses, two-week-old thalli were treated with 25 μM OPDA or wounding, and collected after the indicated time. The RNA extracted with FavorPrep Plant Total RNA Mini Kit (Vienna, Austria), whereas genomic DNA was removed by DNase digestion. Finally, sample quality was evaluated in a Bioanalyzer 2100 (Agilent Technologies, Santa Clara, CA, USA). Three independent biological replicates per sample were analysed. For quantitative PCR (qPCR) analysis, expression of the marker genes was analysed by qPCR using Mp*ACT* (*Mp6g11010*) as a reference gene. Experiments were repeated three times with similar results. Significant difference in gene expression was analysed using the one-way ANOVA/Tukey HSD post hoc test (P < 0.05).

For RNA-seq analyses, the Illumina NovaSeq X Plus platform was employed (Novogene, Cambridge UK). Total RNA was extracted and cDNA was synthesize from poly-A. Raw paired-end reads were trimmed using default parameters with fastp tool (Chen et al., 2018). Then, sequences were mapped and quantified to the reference genome MpTak1_v6.1 with salmon (Patro et al., 2017). Differential expression analyses were performed using DESeq2 (Love et al., 2014) following their manual instruction, and final filters of p-value ≤ 0.05 and log2 of Fold Change ≥ 1 and ≤ −1 for up- and down-regulated DEGs, respectively. No fold-change filter was applied to the genotype:treatment interaction effect approach. GO terms enrichment analysis were performed using topGO (Alexa & Rahnenfuhrer, 2024), performing “new” function with biological processes and “runTest” function with “weight01” algorithm and “fisher” statistic. WGCNA (Langfelder & Horvath, 2008) tool was used for co-expression analysis, running the blockwiseModules function with TOMtype = “signed” and power = 16. PCA plots of the GO terms were performed using a custom modification of the “rrgo” package (Sayols, 2023). Scores similarity matrixes between terms were created using the calculateSimMatrix function, with “org.At.tair.db” as database and *simRel* (“Rel”) similarity method. Scores were calculated from p-values obtained previously from topGO. Kmeans was used to cluster the data, using Silhouette method for calculating the optimal number of clusters (centers). All plots of transcriptional data were generated using ggplot2 package (Wickham, 2016), except for venn diagrams, that were generated using VennDiagram package (Chen & Boutros, 2011). #E64B35FF and #4DBBD5FF were selected as colors, selected from “nrc” palette of ggsci package.

#### Herbivory assays

Ten plants per each genotype were grown in plates for 4 weeks. Then, 12 first-instar larvae of *S. exigua* (Entocare, Wageningen, the Netherlands) were released in each plate. Five to ten plates were used for each genotype. After 7 to 8 days of feeding, larvae were individually collected and weighed on a precision balance. The assays were repeated at least three times with similar results. The experiment was repeated four times with similar results. Data are shown as mean ± SD.

## Supplemental Figure legends

**Supplemental Figure S1.** Chemical structure of the putative dn-OPDA conjugates. The structures of the putative dn-*iso*-OPDA to amino acids identified in untargeted or studied by targeted metabolomic analyses are shown. The quantification of these compounds is presented in Figure 1 and S2.

**Supplemental Figure S2.** Accumulation of putative dn-OPDA-aa conjugates after wounding. Accumulation of putative dn-OPDA conjugates with different amino acids in wild-type Tak-1 (WT) plants at indicated time points after wounding. Peak areas are reported in arbitrary units (AU). Untreated plants, time 0, were included as control. Data shown as mean ± s.d. of four biological replicates, 3-5 plants per each replicate. Asterisks indicate significant differences between wound-induced accumulation of control and wounded plants for each compound according to a t-test (P < 0.05).

**Supplemental Figure S3.** Mp*GH3A* expression is induced by OPDA and wounding in a COI1- and MYC-dependent manner. (A) Analysis of gene expression by microarray study of Mp*GH3* in mock and in response to OPDA (2 hours at 25 μM) in indicated wild-type Tak-1 (WT) and Mp*mycy-1* mutant plants (Peñuelas et al., 2019). (B) qRT-PCR analysis of Mp*GH3A* expression in WT and Mp*coi1-2* mutant plants after wounding. Untreated plants (0) were included as control. Plant tissue was collected after the indicated time. Expression of Mp*GH3A* was normalized against Mp*ACT*. Data shown as mean ± s.d. of three biological replicates, 3-5 plants per each replicate. Experiments were repeated three times with similar results. Letters indicate significant different samples according to the one-way ANOVA/Tukey HSD post hoc test (P < 0.05).

**Supplemental Figure S4.** Generation and analyses of Mp*gh3a^ge^* mutant alleles. (A) Scheme of Mp*GH3A* gene of Marchantia. Grey blocks represent exons and dark grey block highlights the sequence encoding for the GH3 domain. Dashed lines represent deletions in Mp*gh3a^ge^*alleles; frame-shifted sequences in Mp*gh3a^ge^* alleles causing premature stop codons (highlighted in red) are shown. (B) Accumulation of dn-*cis*-OPDA in wild-type Tak-1 (WT) and Mp*gh3a^ge^* mutant plants at indicated time points after mechanical wounding. The experiment was repeated three times with similar results. Data shown as mean ± s.d. of four biological replicates, 3-5 plants per each replicate. Letters indicate significant different samples according to the one-way ANOVA/Tukey HSD post hoc test (P < 0.05).

**Supplemental Figure S5.** Expression of dn-OPDA marker genes in Mp*gh3a^ge^* mutant plants. qRT-PCR analysis of OPDA-regulated genes in wild-type Tak-1 (WT), Mp*coi1-2*, Mp*gh3a-1^ge^* and Mp*gh3a-2^ge^* plants in response to wounding (A-B). Untreated plants (0) were included as control (A). Plants were collected after the indicated times (B). Expression of the Mp*BHLH4* (Mp*2g00930*) and Mp*DIR* (Mp*5g16510)*) genes was normalized against Mp*ACT* (Mp*6g11010*). Data shown as mean ± s.d. of three biological replicates, 3 plants per each replicate. Experiments were repeated three times with similar results. Letters indicate significant different samples according to the one-way ANOVA/Tukey HSD post hoc test (P < 0.05).

**Supplemental Figure S6.** Transcriptomic analysis of *Mpgh3a-1^ge^* in basal conditions. (A) Principal Component Analysis (PCA) plot of the 1000 top genes. (B) Venn diagram of enriched Gene Ontology (GO) terms representing up-regulated DEGs after dn-OPDA treatment in Tak-1 (red) and GO terms representing up-regulated DEGs in *Mpgh3a-1^ge^*compared to Tak-1 in Mock conditions (blue). (C) PCA plot of Gene Ontology (GO) terms based on their score, representing up-regulated DEGs in Tak-1 after dn-OPDA treatment compared to Tak1 in Mock conditions. Each cluster is identified with a different shape and delimited by a dashed circle. GO terms induced by OPDA treatment in WT pants are highlighted in red.

**Supplemental Figure S7.** Gene Ontology and WGCNA analyses of *Mpgh3a-1^ge^* in response to OPDA treatment. (A) Venn diagram of differentially expressed genes (DEGs) up-regulated after dn-OPDA treatment in Tak-1 (red) and DEGs from the genotype:treatment interaction (blue). (B) PCA plot of Gene Ontology (GO) terms based on their score, representing 117 DEGs from turquoise module generated by Weighted Gene Co-expression Network Analysis (WGCNA) after using DEGs from the genotype:treatment interaction effect. Each cluster is identified with a different shape and delimited by a dashed circle. GO terms induced by OPDA treatment in WT pants are highlighted in red.

**Supplemental Figure S8.** Synthetic scheme of dinor-*iso*-OPDA-L-Amino acid conjugates. Schematic representation of the chemical reaction for the synthesis of dinor-*iso*-OPDA-L-Amino acid conjugates. Reagents and conditions: (a) ClCO_2_Et, Et_3_N, THF, 0 ℃; L-amino acid, DIPEA, H_2_O.

**Supplemental table S1**. List of genes shown in RNA-seq analyses.

**Supplemental table S2**. List of GO terms shown in RNA-seq analyses.

**Supplemental table S3**. List of primers used in this work.

## Acknowledgments and Funding

W.L. was funded by the PhD CSC fellowship 202210190002. This work was funded by the Spanish Ministry of Science and Innovation grant TED2021-129735B-I00 and PID2022-140766OB-I00 funded by the MCIN/AEI/10.13039/501100011033 and the European Union ‘‘NextGenerationEU’’/PRTR to R.S. and A.C. This work was also funded by Grant-in-Aid for Scientific Research from JSPS (Japan) No. 23H00316 and JPJSBP120239903, and a Grant-in-Aid for Transformative Research Areas (A) “Latent Chemical Space” (JP23H04880 and JP23H04883) from the Ministry of Education, Culture, Sports, Science and Technology (Japan) to M.U.

## Competing interests

None declared.

## Author Contributions

Conceptualization, R.S. and A.C.; Performed research, W.L. A.Z. A.T. J.C.T. T.K.; Data analysis, W.L. A.Z. S.M. J.C.T. M.C. R.S. A.C.; Writing - original draft preparation, R.S. and A.C.; Manuscript review and editing, all authors; Experimental supervision, M.U. M.C. JM.G.M. R.S. and A.C.; Funding acquisition, R.S. and A.C. All authors have read and agreed to the published version of the manuscript.

## References

Abbas, M., Hernández-García, J., Pollmann, S., Samodelov, S. L., Kolb, M., Friml, J., Hammes, U. Z., Zurbriggen, M. D., Blázquez, M. A., Alabadí, D. (2018). Auxin methylation is required for differential growth in Arabidopsis. Proceedings of the National Academy of Sciences of the United States of America, 115: 6864–6869

Alexa, A., Rahnenfuhrer, J. (2024). topGO: Enrichment Analysis for Gene Ontology. R package version 2.56.0.

Bartel B, Fink G. (1995). ILR1, an amidohydrolase that releases active indole-3-acetic acid from conjugates. Science 268: 1745–1748

Blázquez MA, Nelson DC, Weijers D. (2020) Evolution of plant hormone response pathways. Annu Rev Plant Biol. 71: 327–353.

Boutet S., Barreda L., Perreau F., Totozafy J., Mauve C., Gakière B., Delannoy E., Martin-Magniette M., Monti A., Lepiniec L., et al., (2022) Untargeted metabolomic analyses reveal the diversity and plasticity of the specialized metabolome in seeds of different Camelina sativa genotypes Plant J. 1–19

Bowman, J. L., Kohchi, T., Yamato, K. T., Jenkins, J., Shu, S., Ishizaki, K., Yamaoka, S., Nishihama, R., Nakamura, Y., Berger, F., et al. (2017). Insights into Land Plant Evolution Garnered from the Marchantia polymorpha Genome. Cell, 171: 287–304.e15

Bowman JL, Sandoval EF, Kato H (2021) On the evolutionary origins of land plant auxin biology. Cold Spring Harb Perspect Biol 13: a040048

Brunoni, F., Collani, S., Casanova-Sáez, R., Šimura, J., Karady, M., Schmid, M., Ljung, K., Bellini, C. (2020). Conifers exhibit a characteristic inactivation of auxin to maintain tissue homeostasis. New Phytologist, 226: 1753–1765

Brunoni F, Pěnčík A, Žukauskaitė A, Ament A, Kopečná M, Collani S, Kopečný D, Novák O (2023) Amino acid conjugation of oxIAA is a secondary metabolic regulation involved in auxin homeostasis. New Phytol 238: 2264–2270

Brunoni, F., Široká, J., Mik, V., Pospíšil, T., Kralová, M., Ament, A., Pernisová, M., Karady, M., Htitich, M., Ueda, M., et al. (2023). Conjugation of cis-OPDA with amino acids is a conserved pathway affecting cis-OPDA homeostasis upon stress responses. BioRxiv, 2023.07.18.549545

Campanella, J. J., Kurdach, S., Bochis, J., Smalley, J. (2018). Evidence for Exaptation of the Marchantia polymorpha M20D Peptidase MpILR1 into the Tracheophyte Auxin Regulatory Pathway. Plant Physiology, 177: 1595–1604

Casanova-Sáez, R., Mateo-Bonmatí, E., Šimura, J., Pěnčík, A., Novák, O., Staswick, P., Ljung, K. (2022). Inactivation of the entire Arabidopsis group II GH3s confers tolerance to salinity and water deficit. The New Phytologist, 235: 263–275

Chen Y, Shen H, Wang M, Li Q, He Z (2013). Salicyloyl-aspartate synthesized by the acetyl-amido synthetase GH3.5 is a potential activator of plant immunity in Arabidopsis. Acta Biochim Biophys Sin 45: 827–36

Chen, H., Boutros, P.C. (2011) VennDiagram: a package for the generation of highly-customizable Venn and Euler diagrams in R. BMC Bioinformatics, 12, 35 10.1186/1471-2105-12-35

Chen, S., Zhou, Y., Chen Y., Gu, J. (2018). fastp: an ultra-fast all-in-one FASTQ preprocessor, Bioinformatics, 34: 17–1

Chini, A., Fonseca, S., Chico, J. M., Fernández-Calvo, P., Solano, R. (2009). The ZIM domain mediates homo- and heteromeric interactions between Arabidopsis JAZ proteins. Plant Journal, 59: 77–87

Chini, A., Fonseca, S., Fernández, G., Adie, B., Chico, J. M., Lorenzo, O., García-Casado, G., López-Vidriero, I., Lozano, F. M., Ponce, et al. (2007). The JAZ family of repressors is the missing link in jasmonate signalling. Nature, 448: 666–671

Chini A, Gimenez-Ibanez S, Goossens A, Solano R. (2016). Redundancy and specificity in jasmonate signalling. Current Opinion in Plant Biology 33: 147–156.

Chini, A., Monte, I., Zamarre, A. M., García-Mina, J. M., Solano, R. (2023). Evolution of the jasmonate ligands and their biosynthetic pathways. New Phytologist, 238: 2236–2246

Chini, A., Monte, I., Zamarreño, A. M., Hamberg, M., Lassueur, S., Reymond, P., Weiss, S., Stintzi, A., Schaller, A., Porzel, A., et al. (2018). An OPR3-independent pathway uses 4,5-didehydrojasmonate for jasmonate synthesis. Nature Chemical Biology, 14: 171–178

Farmer, E. E., Davoine, C. (2007). Reactive electrophile species. Current Opinion in Plant Biology, 10: 380–386

Fernández-Calvo, P., Chini, A., Fernández-Barbero, G., Chico, J.-M., Gimenez-Ibanez, S., Geerinck, J., Eeckhout, D., Schweizer, F., Godoy, M., Franco-Zorrilla, J. M., et al. (2011). The Arabidopsis bHLH transcription factors MYC3 and MYC4 are targets of JAZ repressors and act additively with MYC2 in the activation of jasmonate responses. The Plant Cell, 23: 701–715

Fonseca, S., Chini, A., Hamberg, M., Adie, B., Porzel, A., Kramell, R., Miersch, O., Wasternack, C., Solano, R. (2009). (+)-7-iso-Jasmonoyl-L-isoleucine is the endogenous bioactive jasmonate. Nature Chemical Biology, 5: 344–350

Fu, W., Jin, G., Jiménez-Alemán, G. H., Wang, X., Song, J., Li, S., Lou, Y., Li, R. (2022). The jasmonic acid-amino acid conjugates JA-Val and JA-Leu are involved in rice resistance to herbivores. Plant, Cell & Environment, 45: 262–272

Gasperini, D., Howe, G. A. (2024). Phytohormones in a universe of regulatory metabolites: lessons from jasmonate. Plant Physiology, 195: 135–154

Gutierrez, L., Mongelard, G., Flokova, K., Pacurar, D. I., Novak, O., Staswick, P., Kowalczyk, M., Pacurar, M., Demailly, H., Geiss, G., & Bellini, C. (2012). Auxin controls Arabidopsis adventitious root initiation by regulating jasmonic acid homeostasis. Plant Cell, 24: 2515–2527

Howe, G. A., Major, I. T., Koo, A. J. (2018). Annual Review of Plant Biology Modularity in Jasmonate Signaling for Multistress Resilience. Annu Rev Plant Biol. 69:387–415

Jez, J. M. (2022). Connecting primary and specialized metabolism: Amino acid conjugation of phytohormones by GRETCHEN HAGEN 3 (GH3) acyl acid amido synthetases. Current Opinion in Plant Biology, 66: 102194

Jimenez Aleman, G. H., Thirumalaikumar, V. P., Jander, G., Fernie, A. R., Skirycz, A. (2022). OPDA, more than just a jasmonate precursor. Phytochemistry, 204: 113432

Kato, H., Kouno, M., Takeda, M., Suzuki, H., Ishizaki, K., Nishihama, R., Kohchi, T. (2017). The Roles of the Sole Activator-Type Auxin Response Factor in Pattern Formation of Marchantia polymorpha. Plant & Cell Physiology, 58: 1642–1651

Katsir, L., Schilmiller, A. L., Staswick, P. E., Sheng, Y. H., Howe, G. A. (2008). COI1 is a critical component of a receptor for jasmonate and the bacterial virulence factor coronatine. Proceedings of the National Academy of Sciences of the United States of America, 105: 7100–7105

Kneeshaw, S., Soriano, G., Monte, I., Hamberg, M., Zamarreño, A. M., Garcıa-Mina, J. M., Franco-Zorrilla, J. M., Kato, N., Ueda, M., Fernanda Rey-Stolle, M., et al. (2022). Ligand diversity contributes to the full activation of the jasmonate pathway in Marchantia polymorpha. Proceedings of the National Academy of Sciences of the United States of America, 119: e2202930119

Kowalczyk, M., Sandberg, G. (2001). Quantitative analysis of indole-3-acetic acid metabolites in arabidopsis. Plant Physiology, 127: 1845–1853

Kubota A, Ishizaki K, Hosaka M, Kohchi T. (2013) Efficient Agrobacteriummediated transformation of the liverwort Marchantia polymorpha using regenerating thalli. Bioscience, Biotechnology, and Biochemistry 77: 167–172

Langfelder, P., Horvath, S., (2008). WGCNA: an R package for weighted correlation network analysis. BMC Bioinformatics, 9: 559

LeClere, S., Tellez, R., Rampey, R. A., Matsuda, S. P. T., Bartel, B. (2002). Characterization of a Family of IAA-Amino Acid Conjugate Hydrolases from Arabidopsis. Journal of Biological Chemistry, 277: 20446–20452

Li, M., Yu, G., Cao, C., Liu, P. (2021). Metabolism, signaling, and transport of jasmonates. Plant Communications, 2: 100231

Love, MI., Huber, W., Anders, S. (2014). Moderated estimation of fold change and dispersion for RNA-seq data with DESeq2. Genome Biology, 15: 550

Monte, I., Caballero, J., Zamarreño, A. M., Fernández-Barbero, G., García-Mina, J. M., Solano, R. (2022). JAZ is essential for ligand specificity of the COI1/JAZ co-receptor. Proceedings of the National Academy of Sciences of the United States of America, 119: e2212155119

Monte, I., Franco-Zorrilla, J. M., García-Casado, G., Zamarreño, A. M., García-Mina, J. M., Nishihama, R., Kohchi, T., & Solano, R. (2019). A Single JAZ Repressor Controls the Jasmonate Pathway in Marchantia polymorpha. Molecular Plant, 12: 185–198

Monte, I., Ishida, S., Zamarreño, A. M., Hamberg, M., Franco-Zorrilla, J. M., García-Casado, G., Gouhier-Darimont, C., Reymond, P., Takahashi, K., García-Mina, J. M., et al. (2018). Ligand-receptor co-evolution shaped the jasmonate pathway in land plants. Nature Chemical Biology, 14: 480–488

Monte, I., Kneeshaw, S., Franco-Zorrilla, J. M., Chini, A., Zamarreño, A. M., García-Mina, J. M., Solano, R. (2020). An Ancient COI1-Independent Function for Reactive Electrophilic Oxylipins in Thermotolerance. Current Biology, 30: 962–971.e3

Pang Z., Chong J., Zhou G., de Lima Morais D. A., Chang L., Barrette M., Gauthier C., Jacques P.E., Li S., Xia J. (2021). MetaboAnalyst 5.0: narrowing the gap between raw spectra and functional insights. Nucleic Acids Res. 49, W388–W396.

Park, S. W., Li, W., Viehhauser, A., He, B., Kim, S., Nilsson, A. K., Andersson, M. X., Kittle, J. D., Ambavaram, M. M. R., Luan, S., Esker, A. R., Tholl, D., Cimini, D., Ellerström, M., Coaker, G., Mitchell, T. K., Pereira, A., Dietz, K. J., Lawrence, C. B. (2013). Cyclophilin 20-3 relays a 12-oxo-phytodienoic acid signal during stress responsive regulation of cellular redox homeostasis. Proceedings of the National Academy of Sciences of the United States of America, 110: 9559–9564

Patro, R., Duggal, G., Love, M.I., Irizarry, R.A., Kingsford, C. (2017). Salmon: fast and bias-aware quantification of transcript expression using dual-phase inference. Nat Methods. 2017 Apr; 14(4): 417–419

Peñuelas, M., Monte, I., Schweizer, F., Vallat, A., Reymond, P., García-Casado, G., Franco-Zorrilla, J. M., Solano, R. (2019). Jasmonate-Related MYC Transcription Factors Are Functionally Conserved in Marchantia polymorpha. The Plant Cell, 31: 2491–2509

Rampey, R. A., LeClere, S., Kowalczyk, M., Ljung, K., ran Sandberg, G., Bartel, B. (2004). A Family of Auxin-Conjugate Hydrolases That Contributes to Free Indole-3-Acetic Acid Levels during Arabidopsis Germination 1. Plant Physiology, 135: 978–988

Sayols, S. (2023). rrvgo: a Bioconductor package to reduce and visualize Gene Ontology terms. microPublication Biology

Routaboul, J.M., Kerhoas, L., Debeaujon, I., Pourcel, L., Caboche, M., Einhorn, J. et al. (2006) Flavonoid diversity and biosynthesis in seed of Arabidopsis thaliana. Planta, 224: 96–107

Sheard, L. B., Tan, X., Mao, H., Withers, J., Ben-Nissan, G., Hinds, T. R., Kobayashi, Y., Hsu, F. F., Sharon, M., Browse, J., et al. (2010). Jasmonate perception by inositol-phosphate-potentiated COI1-JAZ co-receptor. Nature, 468: 400– 405

Široká J, Brunoni F, Pěnčík A, Mik V, Žukauskaitė A, Strnad M, Novák O, Floková K (2022) High-throughput interspecies profiling of acidic plant hormones using miniaturised sample processing. Plant Methods 18: 122

Soriano, G., Kneeshaw, S., Jimenez-Aleman, G., Zamarreño, Á. M., Franco-Zorrilla, J. M., Rey-Stolle, M. F., Barbas, C., García-Mina, J. M., Solano, R. (2022). An evolutionarily ancient fatty acid desaturase is required for the synthesis of hexadecatrienoic acid, which is the main source of the bioactive jasmonate in Marchantia polymorpha. New Phytologist, 233: 1401–1413

Staswick, P. E., Serban, B., Rowe, M., Tiryaki, I., Maldonado, M. T., Maldonado, M. C., Suza, W. (2005). Characterization of an Arabidopsis enzyme family that conjugates amino acids to indole-3-acetic acid. Plant Cell, 17: 616–627.

Staswick, P. E., Tiryaki, I. (2004). The oxylipin signal jasmonic acid is activated by an enzyme that conjugates it to isoleucine in Arabidopsis. Plant Cell, 16: 2117–2127.

Stepanova, A. N., Alonso, J. M. (2016). Auxin catabolism unplugged: Role of IAA oxidation in auxin homeostasis. Proceedings of the National Academy of Sciences of the United States of America, 113: 10742

Suzuki, H., Kato, H., Iwano, M., Nishihama, R., & Kohchi, T. (2022). Auxin signaling is essential for organogenesis but not for cell survival in the liverwort Marchantia polymorpha. The Plant Cell, 35: 1058–1075. doi10.1093/plcell/koac367

Sztein AE, Cohen JD, Garcia de la Fuente I, Cooke TJ. (1999) Auxin metabolism in mosses and liverworts. Am J Bot. 86:1544–1555

Thines, B., Katsir, L., Melotto, M., Niu, Y., Mandaokar, A., Liu, G., Nomura, K., He, S. Y., Howe, G. A., Browse, J. (2007). JAZ repressor proteins are targets of the SCF(COI1) complex during jasmonate signalling. Nature, 448: 661–665

Wang J., Sakurai H., Kato N., Kaji T., Ueda M. (2021) Syntheses of dinor-*cis/iso*-12-oxo-phytodienoic acid (dn-*cis/iso*-OPDAs), Ancestral Jasmonate Phytohormones of the Bryophyte *Marchantia Polymorpha* L., and their Catabolites. Sci. Rep. 11, 2033.

Wasternack, C., Feussner, I. (2018). The Oxylipin Pathways: Biochemistry and Function. Annu Rev Plant Biol, 69: 363–386

Wasternack, C., Song, S., Napier, R. (2016). Jasmonates: biosynthesis, metabolism, and signaling by proteins activating and repressing transcription. J Exp Bot 69: 1303– 1321

Wasternack, C., Strnad, M. (2018). Jasmonates: News on Occurrence, Biosynthesis, Metabolism and Action of an Ancient Group of Signaling Compounds. International Journal of Molecular Sciences, 19: 2539

Westfall, C. S., Zubieta, C., Herrmann, J., Kapp, U., Nanao, M. H., Jez, J. M. (2012). Structural Basis for PrereceptorModulation of Plant Hormonesby GH3 Proteins. Science, 336: 1708–1711

Wickham, H., (2016). ggplot2: Elegant Graphics for Data Analysis. Springer-Verlag New York.

Xie, D. X., Feys, B. F., James, S., Nieto-Rostro, M., Turner, J. G. (1998). COI1: an Arabidopsis gene required for jasmonate-regulated defense and fertility. Science, 280: 1091–1094.

Yan, J., Li, S., Gu, M., Yao, R., Li, Y., Chen, J., Yang, M., Tong, J., Xiao, L., Nan, F., Xie, D. (2016). Endogenous Bioactive Jasmonate Is Composed of a Set of (+)-7-iso-JA-Amino Acid Conjugates. Plant Physiology, 172: 2154–2164

Yi R, Li Y, Shan X (2024) OPDA/dn-OPDA actions: biosynthesis, metabolism, and signaling. Plant Cell Reports 43: 206

Záveská Drábková L, Dobrev PI, Motyka V. (2015). Phytohormone profiling across the Bryophytes. PLoS ONE 10: e0125411

